# Canonical Analysis of Fluorescent Timer-Anchored Transcriptomes Resolves Joint Temporal and Developmental Progression

**DOI:** 10.64898/2026.03.16.711988

**Authors:** Nobuko Irie, Omnia Reda, Yorifumi Satou, Masahiro Ono

**Author notes:** Correspondence: Dr Masahiro Ono Address: Imperial College London.

## Abstract

Cell-state transitions during development are shaped by multiple concurrent processes, yet single-cell transcriptomic analyses often rely on static molecular profiles or infer time without an experimental anchor. We developed mCanonicalTockySeq, a systems-level framework that reconstructs temporally resolved developmental state spaces by combining signalling history with single-cell RNA sequencing. Using the Nr4a3-Tocky Fluorescent Timer system in developing thymic T cells, we combined scRNA-seq with a molecular clock of strong T-cell receptor signalling to establish an experimentally anchored temporal reference. Using Timer-defined landmark populations, mCanonicalTockySeq constructs a shared state space in which temporal progression and developmental maturation are jointly represented, identifying a Tocky-defined temporal manifold while resolving progression toward CD4 and CD8 single-positive states. This framework recovered biologically coherent dynamics of immediate TCR-response genes, lineage-associated regulators, and agonist-selection-associated programmes. We then extended the framework across species by translating human thymic single-cell transcriptomes into one-to-one mouse ortholog space and projecting them into the mouse Nr4a3-Tocky reference. The projected human cells occupied an interpretable temporal-developmental geometry, and the inferred Tocky-equivalent temporal coordinate showed a significant donor-level association with chronological age. Together, these results establish mCanonicalTockySeq as a general framework for modelling how signalling history and developmental progression are jointly organised in single-cell state space, and illustrate how experimentally anchored reference systems can support comparative analyses across species.

## Introduction

Understanding complex developing systems requires resolving the temporal dimension of cellular state transitions. During development, cell identity is shaped not only by a cell’s observed phenotypic state, but also by the timing, duration, and sequence of instructive signals it has historically experienced ^1^. While single-cell RNA sequencing (scRNA-seq) has revolutionized our ability to map high-dimensional cellular heterogeneity, it remains fundamentally limited by its nature as a destructive, cross-sectional snapshot ^2^. As a result, integrating an anchored, continuous temporal dimension into scRNA-seq data remains a major challenge.

Current computational methods, including pseudotime ^3^ and RNA velocity ^4^, attempt to reconstruct cellular progression from transcriptomic similarity or inferred transcriptional kinetics. While these approaches have been informative, they derive temporal structure from the transcriptome itself rather than integrating an independent experimental record of biological time ^5^. This limitation becomes particularly acute in developing systems, where temporal progression is often confounded by large-scale transcriptomic variation driven by concurrent differentiation. In such settings, cells do not simply move through time or through development alone; rather, temporal progression and developmental maturation intersect within overlapping transcriptional space ^6^.

The Timer-of-cell-kinetics-and-activity (Tocky) system offers an experimental route to address this problem by coupling biological signal induction to a fluorescent molecular clock ^7^. Through the predictable maturation of a Fluorescent Timer protein ^8^, Tocky provides physical temporal landmarks that anchor single-cell transcriptomes to an endogenous record of signalling history ^7^. We previously developed the Nr4a3-Tocky system as an Nr4a3-Timer reporter for analysing T-cell receptor (TCR) signalling dynamics, after identifying Nr4a3 as a TCR-induced gene by applying our Canonical Correspondence Analysis method ^9^ to cross-tissue mouse peripheral and thymic transcriptomic datasets ^8^. However, applying such a temporally anchored system to dynamic developmental trajectories introduces a major analytical challenge: the Tocky-derived temporal readout must be interpreted in the context of the substantial transcriptomic variation generated by the developmental process itself. Previous computational frameworks for Tocky data were primarily designed to extract temporal progression as an isolated structural component ^10–13^, and therefore are less well suited to systems in which time and development proceed simultaneously and share overlapping transcriptional structure.

To address this, we used post-positive selection thymocyte development as a prototypical model of intersecting temporal and developmental axes. Following positive selection, thymocytes undergo large-scale transcriptomic remodelling as they diverge toward CD4 or CD8 single-positive (SP) lineages ^14,15^. Concurrently, these cells are subjected to agonist selection-associated strong TCR signalling, which strongly influences their developmental outcome ^16^. Because strong TCR signalling is superimposed on ongoing lineage maturation, static transcriptomes blur the distinction between developmental progression and transcriptional changes induced by recent signalling events. By applying the Nr4a3-Tocky system to this highly dynamic developmental window, the blue-to-red maturation of the Fluorescent Timer provides a continuous endogenous record of strong TCR signalling over hours to days ^7^. This establishes an experimentally anchored temporal reference in a system where temporal progression and developmental maturation unfold simultaneously.

## Results

### A joint canonical framework resolves thymocyte development across Tocky Time

To investigate thymocyte development in the context of agonist-selection-associated TCR signalling, we combined the Nr4a3-Tocky system with single-cell transcriptomics to construct an experimentally anchored temporal model of post-positive selection thymocytes (**Figure 1a**). In this system, strong TCR signalling induces Nr4a3-driven expression of the Fast-FT Fluorescent Timer protein. Fast-FT spontaneously and irreversibly matures from a blue chromophore to a mCherry-type red chromophore ^8^, with a half-life of approximately 4.1 h, whereas the mature red form is far more stable, with a decay timescale of approximately 122 h ^17^. Consequently, each cell contains Timer proteins spanning multiple maturation states, and the composite blue/red fluorescence profile reports the dynamics of recent, ongoing, and prior transcriptional activity. Tocky therefore resolves signalling dynamics, including both recent immediate events and day-long signalling history, from the composite Timer fluorescence profile of individual cells.

**Figure 1.**
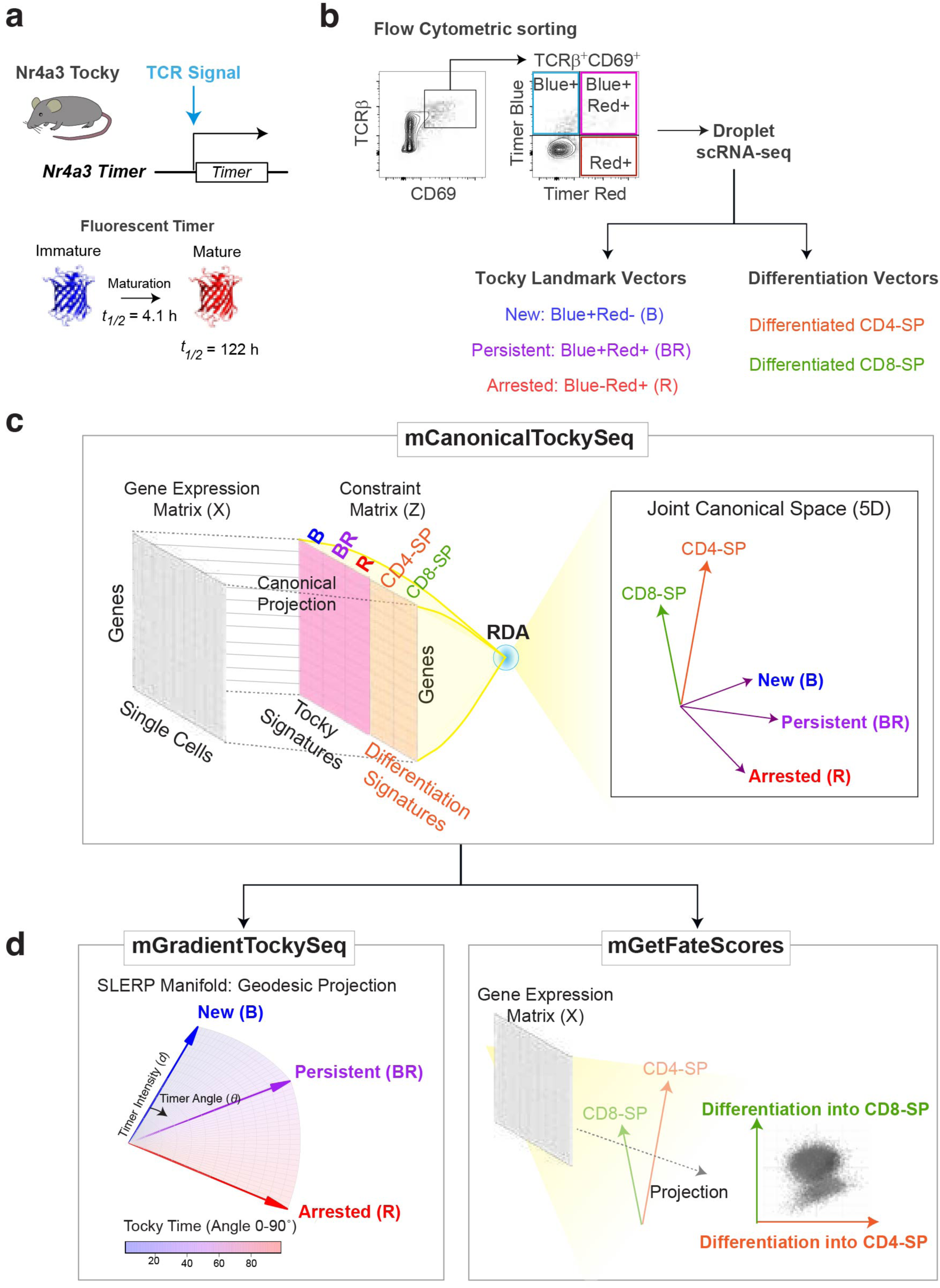
Overview of the mCanonicalTockySeq framework for analysing thymocyte development across Tocky Time. **(a)** Schematic of the Nr4a3-Tocky system. Strong T-cell receptor (TCR) signalling induces Nr4a3-driven Fluorescent Timer expression. The newly translated Timer protein initially emits unstable blue fluorescence, which matures into a stable red fluorescent form. **(b)** Generation of biological landmark populations for canonical modelling. Nr4a3-Tocky post-positive selection CD69^+^TCRβ^+^ thymocytes were flow-sorted into three Timer-defined populations: Blue^+^Red^−^ (New, B), Blue^+^Red^+^ (Persistent, BR), and Blue^−^Red^+^ (Arrested, R), followed by droplet-based single-cell RNA-sequencing. These Timer-defined populations were used to derive Tocky landmark vectors. Developmental endpoint populations, such as differentiated CD4-single positive (SP) and differentiated CD8-SP cells, were used to derive developmental vectors. **(c)** mCanonicalTockySeq constructs a shared canonical space from single-cell gene expression data by constraining the analysis with both Tocky landmark signatures and developmental signatures. Redundancy analysis (RDA) projects the data into a multidimensional canonical Tocky space in which temporal and developmental programmes are represented jointly rather than forced into orthogonal statistical components. **(d)** Biological decomposition by selective readout within the shared canonical space. Left, mGradientTockySeq extracts the temporal component by defining a piecewise SLERP-based Tocky manifold from the New, Persistent, and Arrested landmark vectors. Cells are mapped to this cone-like manifold to obtain Tocky Time (angular position) and Tocky Intensity (radial magnitude). Right, mGetFateScores extracts the developmental component by projecting cells onto developmental endpoint vectors, enabling differentiation into CD4-SP and differentiation into CD8-SP to be quantified in the same canonical space. Schematics are shown to visualise the higher-dimensional framework.

To obtain biological landmarks for this temporal progression, we flow-sorted Nr4a3-Tocky CD69^+^TCRβ^+^ post-positive selection thymocytes into three discrete Timer-defined populations: Blue^+^Red^−^ (New), Blue^+^Red^+^ (Persistent), and Blue^−^Red^+^ (Arrested) (**Figure 1b**). These populations were then profiled by droplet-based single-cell RNA sequencing and used as experimentally defined temporal landmarks.

To analyse how Tocky-defined temporal progression intersects with thymocyte development, we developed *mCanonicalTockySeq*, a computational framework that models temporal progression and developmental maturation in a shared canonical space (**Figure 1c**). In developing thymocytes, Nr4a3-Tocky reports strong TCR signalling associated with agonist selection ^7,12^, whereas developmental progression continues concurrently toward mature CD4 and CD8 states. We therefore reasoned that temporal progression should be analysed within, rather than outside, the broader developmental transcriptomic landscape.

mCanonicalTockySeq constructs a joint canonical space constrained by both Tocky landmarks (New, Persistent, and Arrested) and developmental landmarks (for example, CD4 and CD8 endpoints). Rather than forcing temporal and developmental programmes into orthogonal statistical components, this framework captures them in a shared biologically informed geometry and then decomposes them through selective readout: Tocky landmarks are used to extract temporal progression, whereas developmental landmarks are used to quantify maturation toward lineage endpoints (**Figure 1c**).

Within this shared space, *mGradientTockySeq* extracts the temporal component by using piecewise spherical linear interpolation (SLERP ^18^) to define a continuous Tocky manifold from the New (B), Persistent (BR), and Arrested (R) landmark vectors (**Figure 1d**). Cells are mapped by maximising similarity to interpolated reference vectors along this path, yielding a continuous Tocky Time coordinate, while vector magnitude is quantified separately as Tocky Intensity. As in the previous three-dimensional implementation of *GradientTockySeq* ^19^, this temporal manifold is defined exclusively by the Tocky landmark vectors; however, in the present framework it is extracted within a shared canonical space that also captures developmental structure. This defines a cone-like geometric framework in which temporal progression is encoded by angular position and signal strength by radial distance.

In parallel, *mGetFateScores* extracts the developmental component by projecting cells onto lineage endpoint directions such as CD4 and CD8 in the same canonical geometry (**Figure 1d**). This directional projection yields lineage-oriented fate scores that enable developmental progression to be analysed directly as a function of Tocky Time, revealing how thymocytes mature while traversing Tocky-defined temporal states.

To resolve developmental structure locally across time, we further introduced a trajectory-tube approach (see **Materials and Methods**). Rather than imposing rigid lineage classes, this method identifies gene-centric developmental corridors in fate-score space across overlapping Tocky-time intervals. For each gene of interest, local high-density regions were estimated from expressing cells within each Tocky-time window, and cells repeatedly occupying these regions across overlapping windows were classified as belonging to that gene-associated trajectory tube. This allowed overlapping and transitional developmental identities to be represented explicitly.

Together, these components provide a unified framework for analysing how developmental progression unfolds within experimentally anchored temporal dynamics.

### mCanonicalTockySeq resolves a joint temporal–developmental landscape in Nr4a3-Tocky thymocytes

The temporal landmark libraries were generated from sorted Nr4a3-Tocky populations used directly for scRNA-seq library preparation, rather than being assigned retrospectively by computational classification. We then applied mCanonicalTockySeq to single-cell transcriptomic data from Nr4a3-Tocky thymocytes to test whether the framework could resolve temporal progression and thymocyte development simultaneously in real data. A key intermediate step in this analysis was construction of the shared canonical space itself. Pairwise views of this multidimensional canonical space showed that the Tocky landmark vectors (New, Persistent, and Arrested) and developmental landmark vectors derived from CD4 SP and CD8 SP clusters were embedded jointly in the same geometry, while mGradientTockySeq defined a continuous temporal gradient manifold within that space (**Figure 2a**). Thus, the canonical embedding did not merely reduce dimensionality, but provided the shared biological geometry in which temporal progression and developmental maturation could be resolved simultaneously.

**Figure 2.**
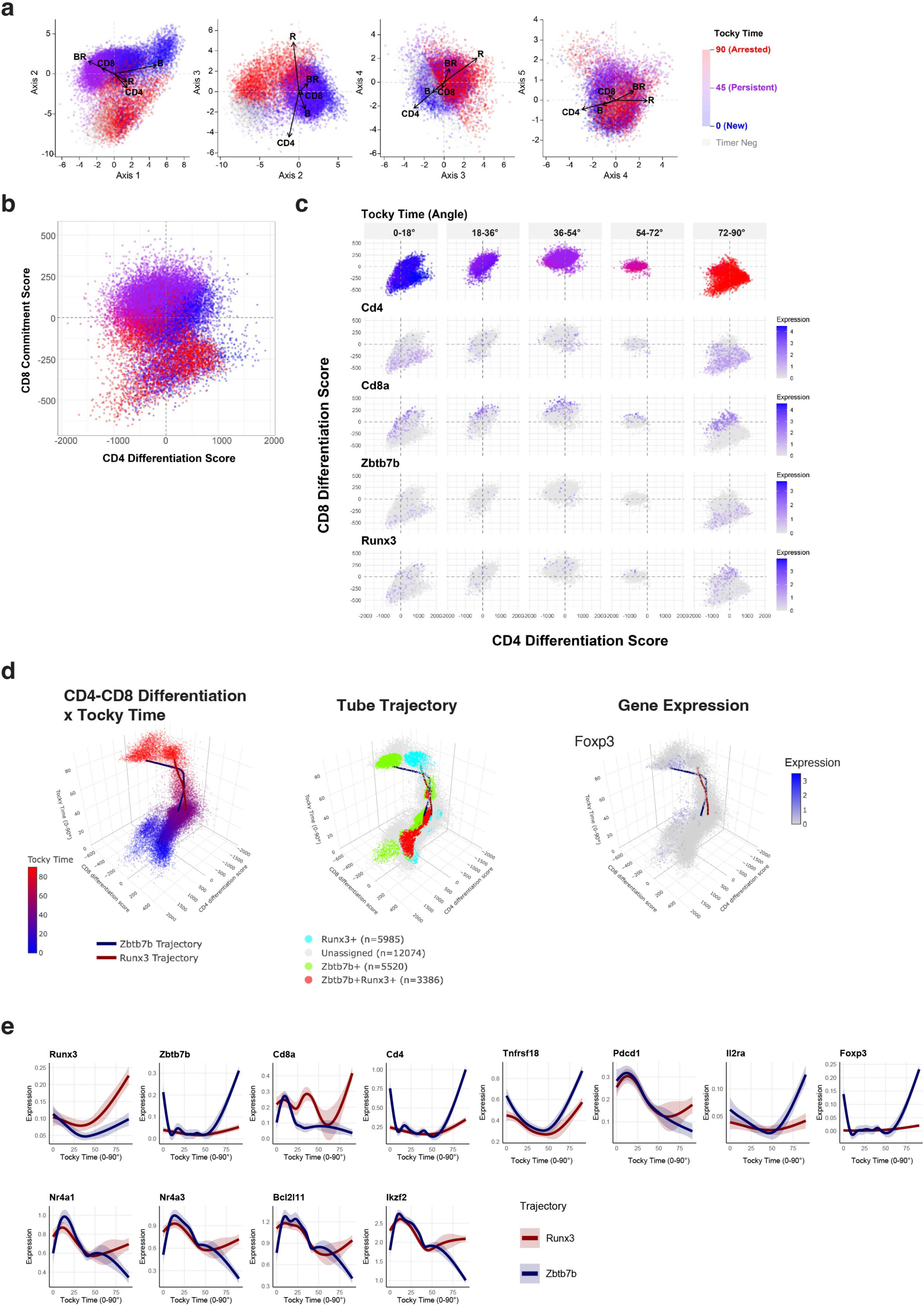
Application of mCanonicalTockySeq to Nr4a3-Tocky thymocytes reveals joint temporal and developmental structure. **(a)** Pairwise views of the multidimensional canonical space generated by mCanonicalTockySeq, coloured by Tocky Time derived from mGradientTockySeq. Arrows indicate the directions of the Tocky landmark vectors (New, Persistent, and Arrested) and developmental landmark vectors (CD4 and CD8), showing that temporal and developmental programmes are embedded jointly in the same canonical geometry. Grey indicates Timer-negative cells. **(b)** Fate-score space derived from mGetFateScores, showing CD4 differentiation score versus CD8 differentiation score, coloured by Tocky Time. **(c)** Gene expression visualised in fate-score space across five equally partitioned Tocky Time intervals (0–18°, 18–36°, 36–54°, 54–72°, and 72–90°). The top row shows the distribution of cells in fate-score space coloured by Tocky Time within each interval. Subsequent rows show expression of Cd4, Cd8a, Foxp3, and Runx3 in the same space. **(d)** Three-dimensional visualisation of the joint developmental-temporal framework using the interactive LaunchTocky3DApp. Left, cells are displayed in CD4 differentiation score × CD8 differentiation score × Tocky Time space, coloured by Tocky Time, with example gene-associated trajectories overlaid. Middle, the same space coloured by overlapping trajectory-tube identities inferred by mExtractTrajectoryTubes. Right, Foxp3 expression displayed in the same three-dimensional space. The overlaid curves are LOESS-smoothed visual summaries of independently inferred gene-associated developmental corridors across Tocky Time. **(e)** Trajectory-wise gene-expression dynamics across Tocky Time. Smoothed GAM expression trends are shown for cells belonging to the Runx3-associated tube (red) or the Zbtb7b-associated tube (blue), with dual-identity cells contributing to each relevant trajectory. Runx3 and Zbtb7b mark the two contrasting developmental programmes, whereas Cd8a and Cd4 report their corresponding lineage-associated states. Tnfrsf18, Pdcd1, Il2ra, and Foxp3 highlight agonist-selection- and regulatory T-cell-associated programmes, and Nr4a1, Nr4a3, Bcl2l11, and Ikzf2 report TCR-signal-associated and downstream developmental responses. Shaded ribbons indicate standard error around the fitted smoothers.

Projection of the same cells into fate-score space using mGetFateScores yielded a two-dimensional developmental representation defined by CD4 differentiation score and CD8 differentiation score (**Figure 2b**). This fate-score space provided a succinct developmental view of the same dataset and showed that thymocytes traversed developmental state space while simultaneously progressing through Tocky-defined temporal states. To examine the organisation of this fate space in greater detail, we visualised selected loci across five equally partitioned Tocky Time intervals, here termed Tocky Loci (0–18°, 18–36°, 36–54°, 54–72°, and 72–90°) (**Figure 2c**). This analysis resolved CD4- and CD8-SP-directed developmental trajectories co-regulated with the lineage-specifying transcription factors Thpok (Zbtb7b) and Runx3, respectively.

We next examined this developmental-temporal structure in three dimensions by plotting cells in CD4 differentiation score × CD8 differentiation score × Tocky Time space (**Figure 2d**). This representation allowed us to ask whether gene-centric analysis based on the two lineage-specifying transcription factors could reveal underlying developmental organisation in a data-oriented manner. To do so, we used a geometric approach to identify cells occupying Runx3-associated or Zbtb7b-associated corridors within the three-dimensional space, designated as *tube trajectories*. In this framework, thymic single cells could be visualised by Tocky Time, by overlapping trajectory-tube identities, or by gene expression. Foxp3 expression provided an illustrative example of how a CD4-SP-associated programme localised within this shared developmental-temporal geometry. Together, these analyses showed that the two tube trajectories captured progressive maturation and developmental divergence, tracing distinct paths that became increasingly separated at later stages of Tocky Time (**Figure 2d**).

Finally, we asked whether these inferred developmental corridors were associated with biologically plausible transcriptional dynamics across Tocky Time. Smoothed GAM trends for cells belonging to the Runx3-associated or Zbtb7b-associated tubes showed distinct trajectory-wise gene-expression dynamics (**Figure 2e**). Immediate TCR-response genes, including Nr4a1 and Nr4a3, were induced early and then declined across Tocky Time, consistent with rapid signalling-dependent activation. In contrast, lineage-associated genes such as Cd4, Cd8a, Runx3, and Zbtb7b showed coordinated but divergent dynamics along the two developmental corridors, consistent with progressive differentiation toward distinct endpoint states. Genes associated with agonist selection, negative selection, or regulatory T-cell-associated programmes also showed distinct temporal profiles within these inferred corridors. Pdcd1 was induced early and then declined, whereas Tnfrsf18, Il2ra, and Foxp3 showed late selective upregulation, particularly along the Zbtb7b-associated trajectory. By contrast, Bcl2l11 and Ikzf2 were induced early in both corridors but diverged later, with more sustained or re-emergent expression along the Runx3-associated path.

Together, these results indicate that the joint canonical framework captures developmental structure without losing the experimentally anchored temporal information carried by the Tocky system, thereby resolving temporally ordered developmental progression together with interpretable gene-expression dynamics in Nr4a3-Tocky thymocytes.

### Cross-species projection of human thymocytes into the Nr4a3-Tocky landmark space reveals Tocky-equivalent progression in developing human T cells

To test whether the Nr4a3-Tocky framework could be extended across species, we projected human thymic single-cell transcriptomes, which profiled dissociated human thymic cells across prenatal, paediatric, and adult stages, including 15 embryonic/foetal thymi spanning 7–17 post-conception weeks and 9 postnatal thymic samples ^20^, into the mouse mCanonicalTockySeq reference space. Human genes were first translated into one-to-one mouse ortholog space and the resulting human thymocyte transcriptomes were then projected using the mouse landmark constraint matrix, thereby positioning human cells within the same canonical geometry defined by the mouse Tocky and developmental landmarks (see **Materials and Methods**, **Figure 3a**). Importantly, the human dataset was not used to re-learn the temporal-developmental model de novo; rather, the experimentally anchored structure learned from the mouse Nr4a3-Tocky system was transferred to the human data as a biological reference coordinate system. This data-learned knowledge transfer enabled temporal and developmental structure in human thymocytes to be interrogated relative to an experimentally grounded mouse model.

**Figure 3.**
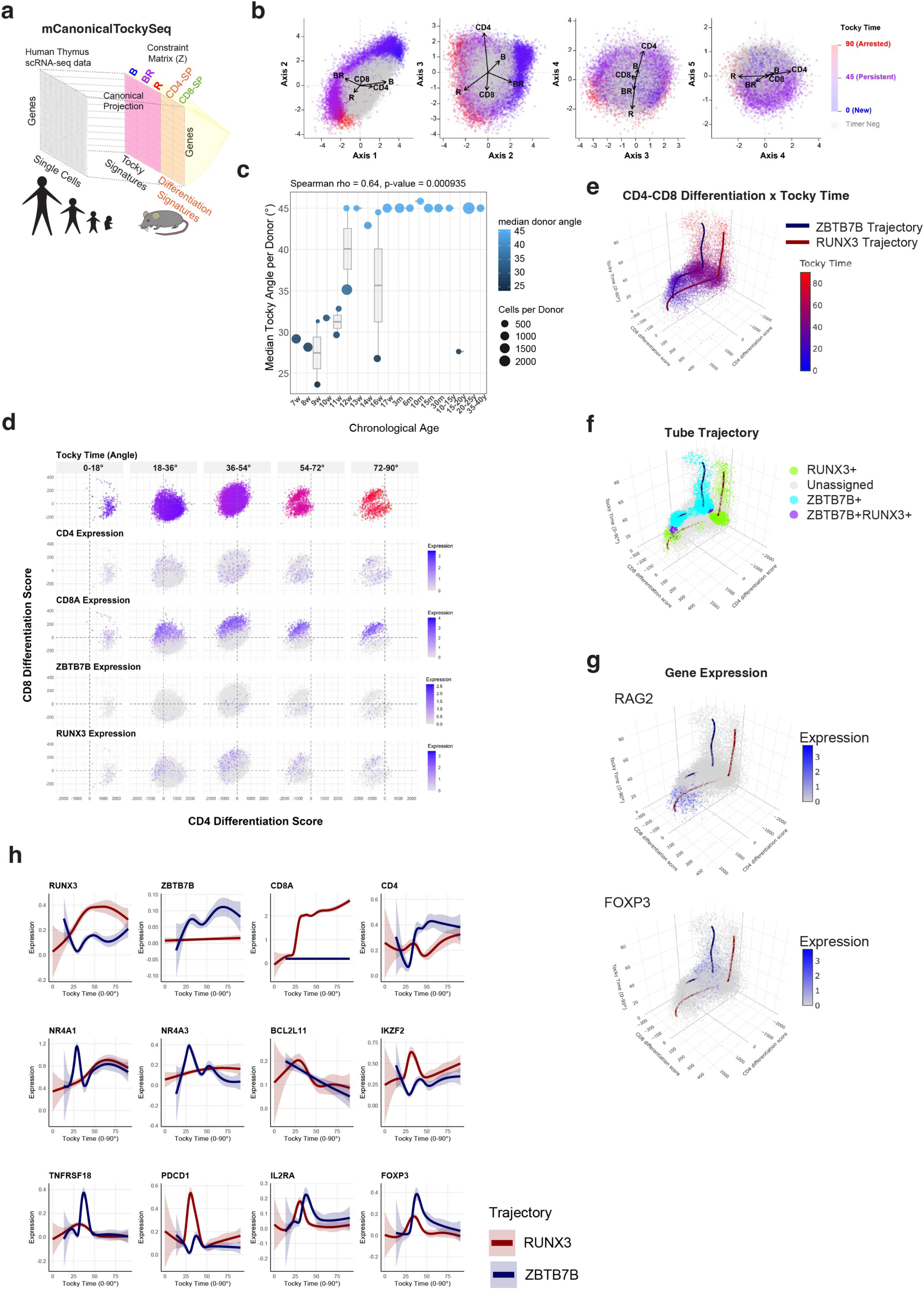
Cross-species projection of human thymocytes into the Nr4a3-Tocky landmark space reveals Tocky-equivalent progression in developing human T cells. **(a)** Schematic of the cross-species projection strategy. Human thymus scRNA-seq data were translated into one-to-one mouse ortholog space and projected into the mouse mCanonicalTockySeq reference using the mouse landmark constraint matrix, thereby positioning human cells within the mouse-defined canonical Tocky space. **(b)** Pairwise views of the projected human cells in the multidimensional canonical space, coloured by Tocky Time inferred by mGradientTockySeq. Arrows indicate the directions of the inherited Tocky landmark vectors (New, Persistent, and Arrested) and developmental landmark vectors (CD4 and CD8), showing that projected human thymocytes occupy the same joint temporal–developmental geometry defined by the mouse reference. Grey indicates Timer-negative cells. **(c)** Donor-level validation of the projected temporal coordinate. For each donor, the median projected Tocky angle was calculated and plotted against ordered chronological age, with point size indicating the number of cells contributed by each donor and point colour indicating median donor angle. Boxplots show the distribution of donor medians within each age group. **(d)** Fate-score space of projected human thymocytes across five equally partitioned Tocky Time intervals (0–18°, 18–36°, 36–54°, 54–72°, and 72–90°). The top row shows the distribution of cells in fate-score space coloured by Tocky Time within each interval. Subsequent rows show expression of CD4, CD8A, ZBTB7B, and RUNX3 in the same space. **(e)** Three-dimensional representation of projected human thymocytes in CD4 differentiation score × CD8 differentiation score × Tocky Time space, coloured by Tocky Time, with example gene-associated developmental trajectories overlaid. **(f)** The same three-dimensional space coloured by overlapping trajectory-tube identities, showing RUNX3-associated, ZBTB7B-associated, dual-identity, and unassigned cells. **(g)** Gene expression displayed in the same three-dimensional developmental–temporal space for RAG2 (top) and FOXP3 (bottom). **(h)** Trajectory-wise gene-expression dynamics across projected Tocky Time. Smoothed GAM expression trends are shown for cells belonging to the RUNX3-associated tube (red) or the ZBTB7B-associated tube (blue), with dual-identity cells contributing to each relevant trajectory. RUNX3 and ZBTB7B mark the two contrasting developmental programmes, whereas CD8A and CD4 report their corresponding lineage-associated states. NR4A1, NR4A3, BCL2L11, and IKZF2 report TCR-signal-associated and downstream developmental responses, whereas TNFRSF18, PDCD1, IL2RA, and FOXP3 highlight agonist-selection- and regulatory T-cell-associated programmes. Shaded ribbons indicate standard error around the fitted smoothers.

Pairwise views of the projected human cells showed that they occupied an interpretable joint canonical space structured by the inherited Tocky landmark vectors (New, Persistent, and Arrested) together with the developmental landmark vectors (CD4 and CD8) (**Figure 3b**). Application of mGradientTockySeq to the projected human cells yielded a continuous Tocky-equivalent temporal coordinate, indicating that the human data could be organised along a progression analogous to the Tocky-defined manifold established in mouse. Thus, the cross-species projection preserved both temporal and developmental structure in a common reference geometry.

We next asked whether this projected temporal coordinate captured biologically meaningful variation in the human dataset. To address this, we related the inferred Tocky angle to donor age across the human thymus samples. At the donor level, the median projected Tocky angle showed a significant positive monotonic association with chronological age (Spearman rho = 0.64, p = 0.000935; **Figure 3c**). Notably, the earliest foetal samples occupied lower projected Tocky-Time states, consistent with the predominance of immature thymocytes in early human thymus, where differentiating αβ T cells are still sparse and progression through DP and SP stages becomes more evident only later in development ^20^. This result thus validated the transferred temporal metric, indicating that the projected human Tocky coordinate reflected biologically coherent developmental structure across human life stages.

Projection of the projected human cells into fate-score space using mGetFateScores further showed that developmental organisation was retained in the transferred reference system (**Figure 3d**). This fate-score space provided a succinct developmental view of the projected human dataset and showed that human thymocytes traversed developmental state space while simultaneously progressing through Tocky-equivalent temporal states. To examine the organisation of this fate space in greater detail, we visualised single cells across five equally partitioned Tocky Time intervals (**Figure 3d**). This analysis resolved CD4- and CD8-directed developmental trajectories co-regulated with the lineage-specifying transcription factors ZBTB7B and RUNX3, respectively, indicating that the mouse-derived canonical reference could be used not only to infer a Tocky-equivalent temporal coordinate in human cells, but also to resolve human developmental progression in parallel.

We next examined this developmental-temporal structure in three dimensions by plotting projected human cells in CD4 differentiation score × CD8 differentiation score × Tocky Time space (**Figure 3e**). Using the geometric approach as in Figure 2, we identified cells occupying RUNX3-associated or ZBTB7B-associated tube trajectories within the three-dimensional space (**Figure 3f**). Expression of genes such as RAG2 and FOXP3 further confirmed the biologically plausible structure of the geometric model, with RAG2 expression confined to early immature thymocytes and Foxp3 found in the ZBTB7B corridor, which is for CD4-SP (**Figure 3g**). Together, these analyses showed that the projected human cells were not only positioned within the mouse-derived reference space, but also supported coherent gene-associated developmental structure within that space.

Finally, we asked whether these inferred developmental corridors in human thymocytes were associated with biologically plausible transcriptional dynamics across projected Tocky Time. Smoothed GAM trends for cells belonging to the RUNX3-associated or ZBTB7B-associated tubes showed distinct trajectory-wise gene-expression dynamics (**Figure 3h**).

RUNX3 and ZBTB7B themselves marked the two contrasting developmental programmes, whereas CD8A and CD4 showed corresponding lineage-associated dynamics along the two projected corridors. Immediate TCR-response and downstream genes also showed corridor-specific temporal behaviour: NR4A1 and NR4A3 displayed prominent early-to-intermediate peaks, whereas BCL2L11 declined progressively and IKZF2 showed a more sustained increase along the RUNX3-associated trajectory. Agonist-selection- and regulatory-associated genes likewise exhibited distinct temporal patterns rather than uniform induction. TNFRSF18 and FOXP3 showed sharp transient peaks within the ZBTB7B-associated corridor, IL2RA peaked in both trajectories but remained more sustained in the ZBTB7B-associated path, and PDCD1 displayed a striking intermediate peak in the RUNX3-associated corridor. These dynamics indicate that projection into the mouse-derived canonical reference did not erase the internal structure of the human dataset. Rather, the transferred framework preserved sufficient structure for coherent, human-specific gene co-regulation to emerge within the shared temporal-developmental geometry.

Collectively, these results support the biological plausibility of the transferred framework and indicate that the mouse Nr4a3-Tocky reference can be used to uncover Tocky-equivalent temporal progression together with interpretable developmental programmes in human thymocytes.

## Discussion

A key implication of the mCanonicalTockySeq framework is that it enables thymocyte development to be analysed without first committing to a rigid classical model of how CD4 and CD8 single-positive (SP) cells emerge ^21^. The framework instead places Tocky-defined temporal progression and movement toward CD4- and CD8-directed states within a common canonical geometry, making it possible to interpret signalling history and developmental fate as coordinated features of the same structure. In this setting, lineage-associated regulators such as Runx3 ^22,23^ and Thpok/Zbtb7b ^24^ serve as gene-centric anchors for identifying modules co-regulated across Tocky Time and developmental state space. Trajectory-tube analysis further represents this organisation as time-local, gene-centric corridors rather than fixed lineage classes. Taken together, these features provide a data-oriented and hypothesis-transparent alternative to more assumption-heavy trajectory interpretations, whose inferred structures can vary substantially depending on the underlying method ^25^.

A further implication of this study is that mCanonicalTockySeq enables temporal-developmental structures learned in experimentally tractable systems to be transferred to settings where direct temporal anchoring is unavailable. Using the mouse Nr4a3-Tocky system as a canonical reference, human thymocytes were projected into the same reference framework without de novo model estimation. The approach therefore operates as an interpretable form of transfer learning ^26^, guided by biological landmarks and oblique canonical geometry rather than black-box optimisation ^27,28^. Crucially, this transfer does not impose a rigid coordinate system on the target data, but instead uses a landmark-defined reference structure that preserves the internal organisation of the human cells while allowing them to be interpreted relative to a Tocky-informed temporal-developmental model. The coherence of the projected human dataset supports this strategy: cells retained an interpretable temporal-developmental manifold, inferred Tocky-equivalent time was significantly associated with chronological donor age, and gene-centric corridors around RUNX3 and ZBTB7B were preserved alongside human-specific co-regulation patterns.

At the same time, the cross-species projection suggests that mouse and human thymocytes share a common temporal-developmental grammar while also raising specific questions about species differences in developmental timing. In both systems, Runx3- and Zbtb7b/Thpok-associated corridors opposed one another and were accompanied by coherent dynamics of lineage-associated, immediate TCR-response, and agonist/Treg-related genes. However, the human projection was consistent with earlier or more transient DP-stage induction of FOXP3 ^29^, IL2RA, and strong agonist-response modules such as NR4A1/NR4A3, PDCD1, and BCL2L11 ^30^, whereas canonical mouse models remain more strongly centred on CD4SP precursor stages ^7,31^. We do not interpret this as a definitive cross-species fate model. Rather, it identifies an open question generated by the projection framework: experimentally anchored temporal reference transfer can reveal both conserved developmental organisation and candidate differences in timing, thereby providing a basis for future mechanistic investigation.

More broadly, the current framework suggests that experimentally anchored temporal-developmental modelling can be extended in both experimental and computational directions. Experimentally, new Tocky systems could be developed for biological settings beyond the thymus. For example, reporters linked to developmental state-transition regulators in neural differentiation, or to pluripotency-associated factors in natural and induced pluripotent systems, could generate landmark reference datasets for the analysis of temporal progression across additional developmental contexts. Computationally, the framework could be expanded through resampling-based stability analysis ^32,33^ and machine-learning-assisted refinement ^34,35^. Such developments may improve the robustness, learning capacity, and cross-system transferability of canonical temporal-developmental models while preserving their interpretability. More generally, these results support the idea that experimentally anchored reference structures can provide a principled basis for analysing temporal progression in systems where direct temporal observation is difficult.

## Materials and Methods

### Multidimensional canonical modelling of Tocky and developmental landmarks

The mCanonicalTockySeq algorithm constructs a shared canonical state space constrained jointly by temporal and developmental landmark populations. The implementation uses a multivariate redundancy-analysis-like procedure followed by truncated singular value decomposition (SVD), implemented in R using the R packages Seurat ^36^, Matrix ^37^, and irlba ^38^.

Single-cell expression data were extracted from the input Seurat object. Biological landmarks were defined from two classes of reference populations: three Tocky states, New (B), Persistent (BR), and Arrested (R), and developmental endpoint populations, here corresponding to CD4 and CD8 states. Features were defined either from a user-supplied gene set or from the union of the top marker genes identified independently for each temporal and developmental landmark group using Seurat. The response matrix (X) was then constructed from the single-cell expression values of these selected genes, with genes as rows and cells as columns. A reference explanatory matrix (Z) was constructed over the same feature set by calculating the average expression of each selected gene across each biological landmark group using Matrix. Temporal and developmental landmark vectors were therefore represented in a common gene space. For internal implementation, landmark columns were relabelled with standard names while preserving user-facing biological labels in the stored provenance metadata.

The explanatory matrix *Z* was optionally scaled column-wise without centring, and the response matrix *X* was standardized across genes for each cell using the base R function scale. Let S denote the standardized response matrix and Z_!"_the scaled explanatory matrix. Constrained projection was then performed by projecting S onto the subspace spanned by *Z_sc_*. In the implementation, this was computed as

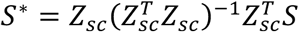

where S^∗^ is the fitted matrix representing the component of the single-cell transcriptomic variation captured by the joint landmark space.

Truncated SVD of S^∗^ was computed using the R package irlba. From this decomposition, the algorithm derived gene-expression scores, fitted cell scores, and cell scores, together with biplot coordinates for the biological landmarks. Cell scores were computed by projecting the standardized single-cell matrix onto the left singular vectors, whereas fitted cell scores were derived from the right singular vectors and singular values. Landmark biplot vectors were obtained by regressing the gene-expression scores onto the unscaled explanatory matrix Z. These outputs were stored in an mCanonicalTockyObj, together with the original expression matrix, explanatory matrix, selected features, metadata, and landmark provenance.

Because temporal and developmental landmarks were modelled jointly, the resulting canonical geometry was not constrained to be orthogonal. Tocky and developmental vectors could therefore overlap, oppose, or intersect according to shared transcriptional structure in the data.

### Human-Mouse cross analysis by projection of new data into a canonical Tocky space using established constraining vectors

Projection of an independent dataset into an existing canonical Tocky reference was performed using mProjectCanonicalTocky. This function reuses the explanatory matrix *Z* stored in a previously generated mCanonicalTockyObj and applies the same constrained projection procedure to a new expression matrix.

The new dataset was first aligned to the reference by intersecting its gene set with the genes present in the stored reference *Z* matrix. In the human–mouse projection analysis, human genes were translated into one-to-one mouse ortholog space before this alignment step. The aligned expression matrix was then standardised, projected onto the fixed reference landmark matrix, and decomposed by truncated SVD using the same sequence of operations described above. In this way, the new cells were placed into the existing canonical coordinate system without modifying the original reference geometry.

The resulting projected object preserved the landmark provenance of the reference model while containing projected gene scores, fitted cell scores, cell scores, and biplot vectors for the new dataset. This projection framework was used to position human thymocytes within the experimentally anchored mouse Nr4a3-Tocky temporal-developmental reference space.

### Temporal mapping with mGradientTockySeq

The mGradientTockySeq algorithm extracts a continuous temporal coordinate from an existing mCanonicalTockyObj by operating specifically on the three Tocky landmark vectors stored in the canonical biplot. The implementation uses a piecewise spherical linear interpolation (SLERP) procedure ^18^ and numerical optimisation in R, using the biplot vectors corresponding to the three Tocky landmarks, New (B), Persistent (BR), and Arrested (R). Developmental vectors present in the same canonical space are not used in this step.

The temporal reference path is defined between the B and BR landmarks and between the BR and R landmarks using piecewise spherical linear interpolation. Unit vectors are first computed for each Tocky landmark, and the angular separation between adjacent landmarks is then calculated. For the two path segments, the angular separations are defined as

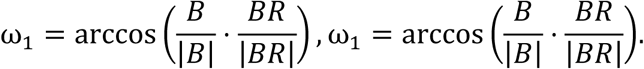

SLERP-derived interpolation weights are then applied to the original landmark vectors to generate continuous reference vectors along each segment. For a local interpolation parameter h ∈ [0,1], the reference vectors are defined as

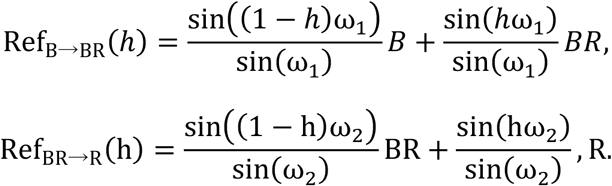

Following constrained projection and truncated singular value decomposition, canonical cell coordinates were computed as

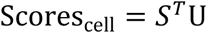

where *S* is the standardized gene-by-cell expression matrix and *U* is the matrix of left singular vectors obtained from the constrained projection matrix. In this representation, each row corresponds to a cell and each column corresponds to a canonical axis. These coordinates define the positions of observed cells in the shared canonical space and were used in all downstream temporal and developmental analyses.

For each cell, projections onto the B, BR, and R landmark vectors are computed. By default, cells with negative projection to all three Tocky landmark vectors are classified as Timer-negative and excluded from temporal assignment. For each remaining cell x_/_, the optimal position along the piecewise reference path is then obtained by numerical optimisation over h ∈ [0,1], maximising similarity between the cell vector and the interpolated reference vector:

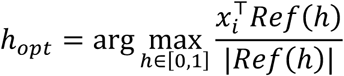

The optimal local position is converted into a global temporal coordinate spanning the full Tocky manifold, and Tocky Time is reported as a scaled angular coordinate from 0 to 90. Canonical intensity is calculated as the Euclidean norm of the cell vector. Expected manifold magnitude at the assigned Tocky position is computed from the corresponding interpolated reference vector, and normalized intensity is defined as the ratio between observed cell magnitude and expected manifold magnitude. An optional quantile-based filter can then be applied to remove cells with low normalised intensity from temporal assignment. In the present study, this filter was used as a biologically motivated enrichment step for cells with sufficient participation in the Tocky-defined manifold, rather than as generic quality control. The resulting TockyData output contains Tocky Time (angle), raw intensity, normalized intensity, and mapping similarity for each cell retained after filtering.

### Developmental projection with mGetFateScores

The mGetFateScores function extracts developmental coordinates by projecting cell coordinates, represented by the canonical cell scores matrix, onto lineage endpoint vectors represented by the corresponding biplot vectors. Developmental fate scores are therefore computed by matrix multiplication of the cell coordinate matrix with the lineage projection matrix. Each cell receives one score for each lineage direction, representing its alignment with the corresponding developmental endpoint within the shared canonical space. In the thymocyte analyses presented here, these developmental endpoints corresponded to CD4 and CD8 states.

### Trajectory-tube extraction as gene-centric developmental corridors across Tocky Time

Trajectory tubes were extracted as local gene-associated structures in fate-score space across Tocky Time. For each cell, the procedure used its Tocky Time (angle) from mGradientTockySeq together with its two lineage-oriented fate scores from mGetFateScores.

For each target gene, overlapping windows of Tocky Time were defined across the interval from 0 to 90 (default window width 10, default step size 1). Within each window, cells with positive expression of the target gene were identified from the Seurat expression matrix. A core expressing subset was then defined as cells with expression greater than or equal to the median of the positive-expression values in that window.

For each window, the local centre of the gene-associated structure was estimated from a two-dimensional kernel density estimate of the core-cell coordinates in fate-score space, and the peak-density location was taken as the local barycentre using the R package MASS ^39^. Local shape was estimated from the covariance matrix of the same core-cell coordinates, with a small diagonal regularisation term added for numerical stability. For all cells in the corresponding Tocky-time window, squared Mahalanobis distance from the local barycentre was calculated, and tube membership was determined relative to the confidence ellipse defined by the chi-square threshold at the selected confidence level (ci).

Because Tocky-time windows overlap, the same cell could be evaluated multiple times for the same gene. Final tube membership was therefore assigned by majority rule across all evaluable windows. Tubes were computed independently for each selected gene, allowing cells to belong to none, one, or multiple tubes simultaneously. Composite identities were then generated by concatenating all positive tube assignments for each cell, with cells lacking any assignment labelled Unassigned.

### Cross-species translation and projection of human thymocytes into the mouse mCanonicalTockySeq reference

For cross-species analysis, human thymic single-cell transcriptomes were translated into mouse ortholog space and then projected into the experimentally anchored mouse mCanonicalTockySeq reference. The human thymus dataset used for cross-species projection was derived from the life-span single-cell atlas reported by Park et al,. which profiled dissociated human thymic cells across prenatal, paediatric, and adult stages, including 15 embryonic/foetal thymi spanning 7–17 post-conception weeks and 9 postnatal thymic samples ^20^. We accessed the corresponding public Human Cell Atlas project (HumanThymicDevelopment; project ID c1810dbc-16d2-45c3-b45e-3e675f88d87b).

Human gene symbols were mapped to mouse ortholog symbols using the R package biomaRt ^40^ through the human Ensembl dataset (hsapiens_gene_ensembl), retrieving the mmusculus_homolog_associated_gene_name attribute. To ensure an unambiguous feature set, only strictly one-to-one human–mouse ortholog mappings were retained, excluding human genes mapping to multiple mouse genes and mouse genes receiving mappings from multiple human genes. The human count and normalized expression matrices were then restricted to the shared ortholog set and relabelled with mouse ortholog symbols. A new Seurat object was constructed from the translated counts, and the translated normalized expression matrix was inserted as the data layer while preserving the original human cell metadata.

To focus the analysis on a developmentally appropriate human thymocyte compartment, the translated human dataset was subset to Rag1-low and Rag2-low cells (Rag1 < 0.5, Rag2 < 0.5). These cells were processed using a standard Seurat workflow for exploratory visualisation and annotation. The translated human expression matrix was then projected into the mouse mCanonicalTockySeq reference using the mProjectCanonicalTocky function. In this step, the landmark constraint matrix *Z* was taken directly from the mouse reference object, and projection was performed only on genes shared between the translated human dataset and the reference *Z* matrix. The same redundancy-analysis-based canonical procedure used for the mouse model was then applied to the translated human data, yielding projected gene scores, cell scores, fitted cell scores, and biplot vectors in the mouse-defined canonical space. The projected human object inherited the landmark provenance of the mouse reference, including the Tocky and developmental endpoint mappings.

Following projection, the human cells were analysed using the same downstream framework as the mouse data. mGradientTockySeq was used to infer a Tocky-equivalent temporal coordinate relative to the mouse-derived New, Persistent, and Arrested landmark vectors, and mGetFateScores was used to derive CD4 and CD8 differentiation scores in the same projected space. Gene-associated developmental corridors were then inferred in the projected human data using mExtractTrajectoryTubes, with ZBTB7B and RUNX3 used as defining gene anchors. In this way, the mouse Nr4a3-Tocky system served as an experimentally anchored biological reference for analysing temporal-developmental structure in human thymocytes.

### Interactive three-dimensional visualisation of developmental-temporal structure

Interactive three-dimensional visualisation was implemented in R using the R package shiny ^41^ for the user interface and the R package plotly ^42^ for three-dimensional rendering. Cells were displayed in CD4 differentiation score × CD8 differentiation score × Tocky Time space using the developmental fate scores, temporal coordinates, and trajectory-tube assignments stored in the analysis object, together with the corresponding Seurat object for retrieval of gene-expression values.

Cells lacking assigned Tocky Time were excluded from the three-dimensional display. To visualise gene-associated trajectories, local centroids were computed independently for each trajectory tube across overlapping Tocky-time windows, and LOESS smoothing was then applied to the resulting coordinates to generate continuous three-dimensional path summaries. These smoothed paths were used as visual representations of gene-associated developmental corridors.

The interactive display supported three colouring modes: Tocky Time, gene-associated identities, and gene expression. In Tocky Time mode, cells were coloured according to their temporal coordinate. In gene-associated identity mode, cells were coloured by composite trajectory-tube assignment, allowing overlapping tube-defined identities to be visualised directly. In gene-expression mode, expression of a selected gene was retrieved from the Seurat expression matrix and displayed on the same three-dimensional coordinate system.

### Mice

The *Nr4a3*-Tocky strain, BAC Tg (*Nr4a3ΔExon3 Fast-FT*) was generated by the Ono group and reported previously ^7^. In this system, the Fluorescent Timer (*Fast-FT* ^8^) gene replaced the first coding exon of the *Nr4a3* gene within the BAC transgene. The mice were bred with *Foxp3*-IRES-GFP (C.Cg-Foxp3tm2Tch/J, Jax ID 006769) and maintained in a C57BL/6 background. All animal procedures were conducted in compliance with the UK Animals (Scientific Procedures) Act 1986, under the oversight of the Animal Welfare and Ethical Review Body (AWERB) at Imperial College London and the Animal Experiment Committee at Kumamoto University.

### Droplet scRNA-seq Library Preparation and Data Preprocessing

Thymic Tocky landmark data were generated by sorting Tocky fractions within CD69^+^TCRβ^+^ population via using a FACSAria (BD Biosciences): New (Blue^+^Red^−^), Persistent (Blue^+^Red^+^), Arrested (Blue^−^Red^+^). Sorted cells were processed immediately for library construction using the Chromium Single Cell 3’ Library & Gel Bead Kit (v3.1, 10x Genomics). Library quality was confirmed by TapeStation (Agilent). High-throughput sequencing was performed by HiSeq X Ten (Illumina) at the Macrogen Japan NGS facility.

## Acknowledgements

This research was supported by KAKENHI research grants from the Japan Society for the Promotion of Science (JSPS) (JP19H05426 and JP24K10259 to MO; JP25K02693 and JP25K22556 to YS), Japan Agency for Medical Research and Development (AMED) (25gm1810001s0104 to MO; JP23wm0325068 and JP25fk0410070 to YS), and JST-ASPIRE Program (JP29jf0126018).

## Author Contributions Statement

MO conceived the study and all computational and mCanonicalTockySeq methodologies, designed joint canonical framework, wrote computational codes, and performed all computational analysis. NI, OR, YS, and MO performed experiments. YS and MO designed experiments and secured funding. MO wrote the manuscript.

## Data Availability

The sequencing data reported in this study have been deposited in the NCBI Sequence Read Archive (SRA) under BioProject accession PRJNA1437868. Interactive three-dimensional visualisations of the Nr4a3-Tocky developmental-temporal models are available at: https://monotockylab.github.io/Nr4a3-Tocky-3D-Models/

## Code Availability

The code reported in this study is publicly available at GitHub: https://github.com/MonoTockyLab/mCanonicalTockySeq

